# Hypo-responsiveness of human alveolar macrophages to IFN-γ is not due to attenuated STAT1 signaling

**DOI:** 10.1101/2022.12.07.519298

**Authors:** Bonnie A Thiel, Kathleen C Lundberg, Daniela Schlatzer, Jessica Jarvela, Qing Li, Rachel Shaw, Sara E Beckloff, Mark R Chance, W Henry Boom, Richard F Silver, Gurkan Bebek

**Affiliations:** Department of Medicine, Case Western Reserve University and University Hospitals Cleveland Medical Center, Cleveland, OH USA; Department of Nutrition, Center for Proteomics and Bioinformatics, Case Western Reserve University School of Medicine, Cleveland, OH USA; Division of Pulmonary, Critical Care, and Sleep Medicine, Louis Stokes Cleveland Department of Veterans Affairs Medical Center, Cleveland, OH USA; Division of Pulmonary, Critical Care, and Sleep Medicine, University Hospitals Case Medical Center and Case Western Reserve University School of Medicine, Cleveland, OH USA; Biobot Analytics, Cambridge, MA USA

**Keywords:** alveolar macrophage, monocyte, IFN-γ, proteome, unlabeled mass spectrometry

## Abstract

Alveolar macrophages (AM) perform a primary defense mechanism in the lung through phagocytosis of inhaled particles and microorganisms. AM are known to be relatively immunosuppressive consistent with the aim to limit alveolar inflammation and maintain effective gas exchange in the face of these constant challenges. How AM respond to T cell derived cytokine signals, which are critical to the defense against inhaled pathogens, is less well understood. For example, successful containment of *Mycobacterium tuberculosis* (Mtb) in lung macrophages is highly dependent on IFN-γ secreted by Th-1 lymphocytes, however, the proteomic IFN-γ response profile in AM remains mostly unknown. In this study, we measured IFN-γ induced protein abundance changes in human AM and autologous blood monocytes (MN). AM cells were activated by IFN-γ stimulation resulting in STAT1 phosphorylation and production of MIG/CXCL9 chemokine. However, the global proteomic response to IFN-γ in AM was dramatically limited in comparison to that of MN (9 AM vs 89 MN differentially abundant proteins). AM hypo-responsiveness was not explained by reduced JAK-STAT1 signaling nor increased SOCS1 expression. These findings suggest that AM have a tightly regulated response to IFN-γ which may prevent excessive pulmonary inflammation but may also provide a niche for the initial survival and growth of Mtb and other intracellular pathogens in the lung.

## INTRODUCTION

To avoid tissue damage due to inflammation, alveolar macrophages (AM) balance the need to respond appropriately to pathogens while limiting local inflammation to preserve the gas-exchange function of the lung. The current paradigm of pulmonary macrophage development suggests that resident AM maintain a localized population with a differentiated immunosuppressive phenotype but can be supplemented with blood monocytes when necessary to respond to an active infection [1]. The activation state of AM when exposed to T cell-derived cytokines has been modeled using monocyte-derived-macrophages but rarely studied in ex-vivo human AM. In the context of *Mycobacterium tuberculosis* (Mtb) infection, IFN-γ stimulates resident lung alveolar macrophages (AM) to produce chemokines, such as MIG (CXCL9) and IP-10 (CXCL10), resulting in recruitment of additional T-cells, blood monocytes (MN) and interstitial macrophages to sites of infection and control of Mtb growth [2,3]. In this comparative study, we sought to clarify the role of differential responses to IFN-γ in the varying inflammatory vs immunosuppressive phenotypes of MN and AM.

Comparative proteomic analysis of whole cell lysates using unlabeled mass spectrometry provides a wide-angle snapshot of the relative abundance of thousands of proteins that can be probed for differences in known functional pathways and networks and suggest novel biomarker targets. In a previous study, we found that the proteomes of human AM and MN overlapped by 70% and there was enrichment of inflammatory response proteins in MN, and protein synthesis/trafficking modules in AM [4]. To follow-up to this baseline study, we modeled T-cell activation of human AM and MN using physiological levels of IFN-γ to compare proteomic responses in cells from healthy Mtb-naïve subjects with the aim of identifying pathways associated with AM immune function in the lung.

## METHODS

### Subject population

Healthy volunteers (non-smokers between 18 and 50 years old) donated blood for monocyte isolation and underwent bronchoscopy with bronchoalveolar lavage (BAL) to obtain AM. Ten donors contributed samples for the exploratory proteomic analysis and 4 additional donors were recruited for validation studies. Subjects were confirmed as Mtb-naïve based on PPD skin testing or IGRA blood tests. Bronchoscopies were performed in the Dahms Clinical Research Unit of University Hospitals Cleveland Medical Center using previously described protocols [5]. All subjects consented to protocols approved by the Institutional Review Board of Case Western Reserve University and the Louis Stokes Cleveland Department of Veterans’ Affairs Medical Center.

### AM and MN isolation

BAL fluid was aliquoted into 50 mL polypropylene tubes and centrifuged at 300 x g for 10 min. Supernatants were removed, and BAL cells were resuspended in Iscove’s Modified Dulbecco’s Medium (IMDM) with 1% penicillin G. BAL cell differentials were determined by light microscopy counting of 300 cells on Wright-Giemsa–stained cytospin preparations (LeukoStat; Fisher Diagnostics, Pittsburgh, PA). The percentage of AM cells among BAL cells was > 90% by light microscopy counting of 300 cells on Wright-Giemsa–stained cytospin preparations (LeukoStat; Fisher Diagnostics, Pittsburgh, PA) and no further purification was done. Given the 90%+ purity, will refer to BAL cells as AM.

Peripheral blood mononuclear cells (PBMC) were isolated from whole blood by Ficoll-Hypaque centrifugation (GE Healthcare) and processed according to manufacturer’s instructions. For proteomic analysis, monocytes were isolated by adherence to polystyrene plates, followed by additional removal of non-adherent lymphocytes via light vortexing of the plates. For validation samples, CD14+CD16-monocytes were isolated from peripheral blood mononuclear cells using immunomagnetic negative selection (EasySep Human Monocyte Isolation Kit, Stem Cell Technologies).

### IFN-γ stimulation

AM and MN cells (1 – 2 × 10^6^) were re-suspended in IMDM with 2% fetal bovine serum and 1% penicillin G, aliquoted into 1.5 ml microfuge tubes, and incubated at 37°C in media with or without 0.5 ng/ml (1.0 unit/ml) human IFN-γ (ThermoFisher Scientific) at time points ranging from 5 minutes to 24 hours. This dose, while considerably lower than that used in many studies of macrophage activation by IFN-γ, was chosen as being physiologically relevant based on assessment of its production in human subjects with latent tuberculosis infection (LTBI) in response to re-exposure to Mtb antigens; further, this dose induced production of IFNγ-inducible chemokines CXCL9 and CXCL10 in BAL samples from both LTBI and Mtb naïve individuals.

### Proteomic analysis sample preparation

Samples were centrifuged at 300 x g for 10 min. Supernatants were harvested for CXCL9 chemokine measurements. Cell pellets for mass spectrometry measures were snap frozen by immersion in liquid nitrogen for 5 min prior to storage at −80°C. When sample collection was complete, cell pellets were thawed and processed simultaneously. 100 μL of 4% SDS with 1X protease inhibitor cocktail (Sigma-Aldrich, St. Louis, MO) was added to cells. Each sample underwent three 5-second pulse sonications with a probe sonicator at 15% amplitude and 5 second rest intervals between repetitions. Lysates were then incubated on ice for 45 min and pulse-sonicated again before an additional 5.5 h incubation on ice.

Following cell lysis, samples were processed using a filter-aided sample preparation (FASP) cleanup protocol with Amicon Ultra MWCO 3K filters (Millipore, Billerica, MA) as previously described (Neufeld 1991 Ann Rev Biochem). Samples were reduced and alkylated on the filters with 10mM dithiothreitol (Acros, Fair Lawn, NJ) and 25mM iodoacetamide (Acros, Fair Lawn, NJ), respectively; after which they were concentrated to a final volume of 40μL in 8M Urea. Protein concentration was performed using the Bradford Method as adapted to the manufacturer’s instructions (Bio-Rad, Hercules, CA).

Five micrograms of total protein were aliquoted for enzymatic digestion. Urea concentration was adjusted to 4M using 50mM Tris pH 8 and proteins were digested with mass spectrometry grade lysyl endopeptidase (Wako Chemicals, Richmond, VA) in an enzyme/substrate ratio of 1:20 for 2 hours at 37°C. Urea concentration was further adjusted to 2M using 50mM Tris pH 8 and Lysyl peptides were additionally digested with sequencing grade trypsin (Promega, Madison, WI) in an enzyme/substrate ratio of 1:20 at 37°C overnight. Finally, samples were diluted in 0.1% formic acid (Thermo Scientific, Rockford, IL) prior to LC-MS/MS analysis. 400fmol of Pierce^®^ Retention Time Calibration Mixture (Thermo Scientific, Rockford, IL) was spiked into samples to track retention times and mass drift across all samples. MN samples showed a maximum retention time drift of 2.07 min whereas AM samples showed a 3.49 min maximum retention time drift across the same 9 tracked peptides. Additionally, the mass drift was accounted for using the same 9 tracked peptides in both cell types; MN showed a maximum 4.15 ppm error whereas AM had a 3.35 ppm maximum mass error.

### Liquid Chromatography and mass spectrometry

For each sample, 400ng of AM and MN peptide digests were loaded on a column in a 14μL injection. Samples were randomized and blanks were added between samples. Resulting data was acquired on an Orbitrap Velos Elite mass spectrometer (Thermo Electron, San Jose, CA) equipped with a Waters nanoAcquity LC system (Waters, Taunton, MA). Peptides were desalted in a trap column (180 μm × 20 mm, packed with C18 Symmetry, 5μm, 100Å, Waters, Taunton, MA) and subsequently resolved in a reversed phase column (75μm x 250 mm nano column, packed with C18 BEH130, 1.7μm, 130Å (Waters, Taunton, MA)). Liquid chromatography was carried out at ambient temperature at a flow rate of 300 nL/min using a gradient mixture of 0.1% formic acid in water (solvent A) and 0.1% formic acid in acetonitrile (solvent B). The gradient employed ranged from 4 to 44% solvent B over 210 min. Peptides eluting from the capillary tip were introduced into the nanospray mode with a capillary voltage of 2.4 kV. A full scan was obtained for eluted peptides in the range of 380–1800 atomic mass units followed by twenty-five data dependent MS/MS scans. MS/MS spectra were generated by collision-induced dissociation of the peptide ions at normalized collision energy of 35% to generate a series of b- and y-ions as major fragments. A one-hour wash was included between each sample.

### Protein identification

Progenesis LC-MS version 4.1 software (Nonlinear Dynamics, Garth Heads, UK) was used to align the retention times (using Progenesis-recommended cut-offs), normalize and identify data features. The IPI-human database from June 2010 (86,392 sequences) was used as a decoy database. FDR was determined by assessment of peptide matches above the identity or homology threshold. Key search parameters were Trypsin + Lys-C for enzyme; a maximum of 2 missed cleavages; peptide charge states of +2 to +4; peptide tolerance of 10 ppm; and MS/MS tolerance of 0.8 Da. Fixed modifications included carbamidomethylation of cysteine residues. Oxidation of methionine was a variable modification.

Tables of all measured peptides including sequences, charge states, modifications and protein identifiers are available in the supplemental tables.

### ELISA

Culture supernatants were collected and stored after IFN-γ stimulation. CXCL9 chemokine concentration was measured using a commercially available ELISA kit (R and D Systems) according to manufacturer’s instructions.

### Western blot analysis

1 × 10^6^ MN and AM cells from each donor were cultured with recombinant human IFN-γ (0.5 ng/ml) overnight. Cells were pelleted and lysed in RIPA buffer (Cell Signaling Technologies), supplemented with Halt Protease and Phosphatase Inhibitor Cocktail (ThermoFisher Scientific), sonicated for 1 minute and centrifuged at 14,000 RPM for 10 minutes at 4°C. Protein concentration in lysates was measured using the Pierce BCA Protein Assay kit (ThermoFisher Scientific) and diluted to a concentration of 1-2 μg/uL. Samples were heated to 95°C for 5 minutes in Laemmli buffer (Alfa Aesar), separated on a Novex 4-20% Tris-glycine gel (ThermoFisher Scientific) and transferred to an Immuno-Blot PVDF membrane (BioRad Laboratories). Membranes were blocked in 5% BSA with 1% Tween 20 for 1 hour then probed overnight at 4°C with the following primary antibodies at a 1:1,000 dilution: Phospho-STAT1 (Cell Signaling Technologies, 9167S, monoclonal), SOCS1 (Cell Signaling Technologies, 3950S, polyclonal), STAT-1 (Cell Signaling Technologies, 14994S, monoclonal), IFNGR1 (Abcam, 134070, monoclonal) and GAPDH (Cell Signaling Technologies, 2118S, monoclonal). The next day, membranes were washed 3 times in TBST (Tris buffered saline with 0.1% Tween20) for 10 minutes and probed for 1 hour at room temperature with the horseradish peroxidase-conjugated mouse anti-rabbit IgG monoclonal antibody at a 1:5,000 dilution (Jackson ImmunoResearch, 211-035-109). Membranes were washed three more times in TBST before protein expression was detected using the Pierce ECL Plus Substrate (ThermoFisher Scientific) on the BioBlot Blue X-Ray film (Laboratory Product Sales). Bands were quantified using ImageJ software (version 1.53a; available from website imagej.nih.gov) and normalized to the GAPDH loading control.

### mRNA measurement by quantitative PCR

Total RNA was purified from the MN and AM (0.5 × 10^6^ cells) using the RNeasy Mini Kit (Qiagen). Total RNA was reverse transcribed using the QuantiTect Reverse Transcription Kit (Qiagen). RT-qPCR samples were run in duplicate on the StepOne Plus Real-Time PCR System (Applied Biosystems) at 95°C for 10 minutes and 60°C for 1 minute, then 40 cycles of 95°C for 15 seconds and 60°C for 1 minute. The total reaction volume was 25 ul. For GAPDH, 12.5 μl of SYBR Green (Roche), 0.75 μl of the forward and reverse primer (OriGene, HP205798), and 10.75 μl of RNase-free water were added to each reaction well followed by 1μl of cDNA. For SOCS1 and STAT-1, 12.5 μl of SYBR Green, 0.75 μl of the forward and reverse primers (OriGene, HP207209 and HP210040) and 9.25 μl of RNase-free water were added to each well followed by 2.5 μl of cDNA.

Primers for Reverse Transcription quantitative PCR:

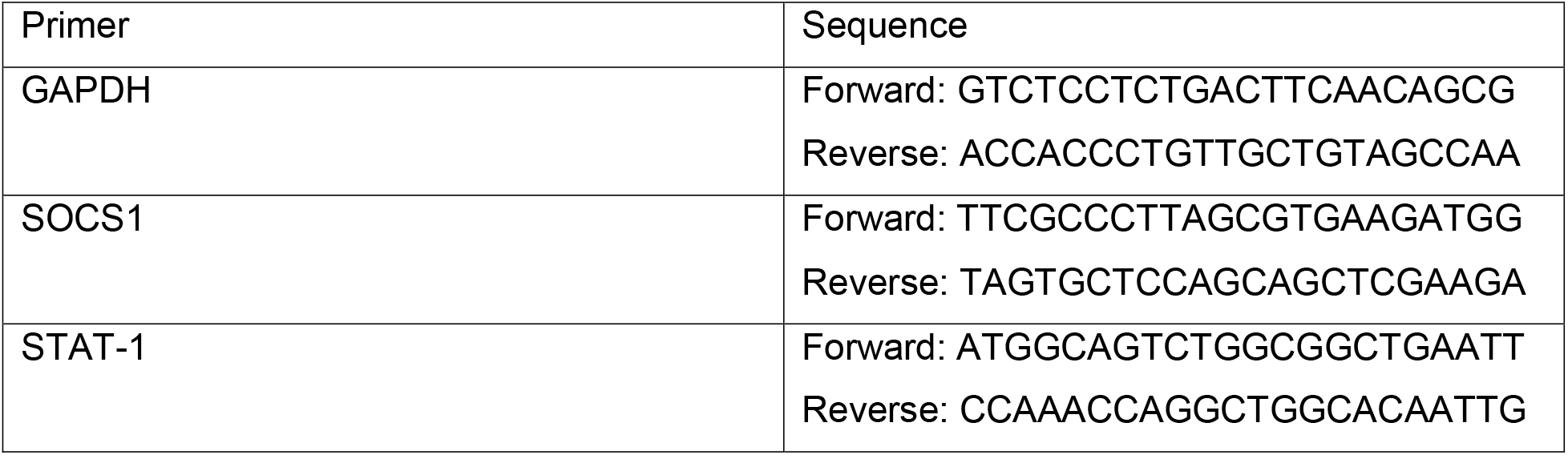

### Data analysis

Data integration and analysis pipeline is shown in figure 1. Data summarization and statistical analyses were done using R (Versions 3.2.3 and 3.5.1, Vienna, Austria). For the proteomic dataset, proteins identified by a single peptide in only one cell type were filtered out of the analysis dataset as these have a high probability of being miss-identified proteins. A linear modeling approach was used to estimate the effect of IFN-γ on protein levels while controlling for the correlated peptide abundance measures within each subject [6,7]. Treatment effect t-statistics, estimated for MN and AM separately, were ranked by the size of the IFN-γ effect and tested for significant enrichment of functionally related proteins using gene sets from the Molecular Signatures Database (MsigDB, version 7.2 accessed November 2020) where the term ‘gene’ refers to a gene-product [8]. The Hallmark set of molecular signatures was used to test for enrichment in the IFN-γ response process [9]

**Figure 1:**
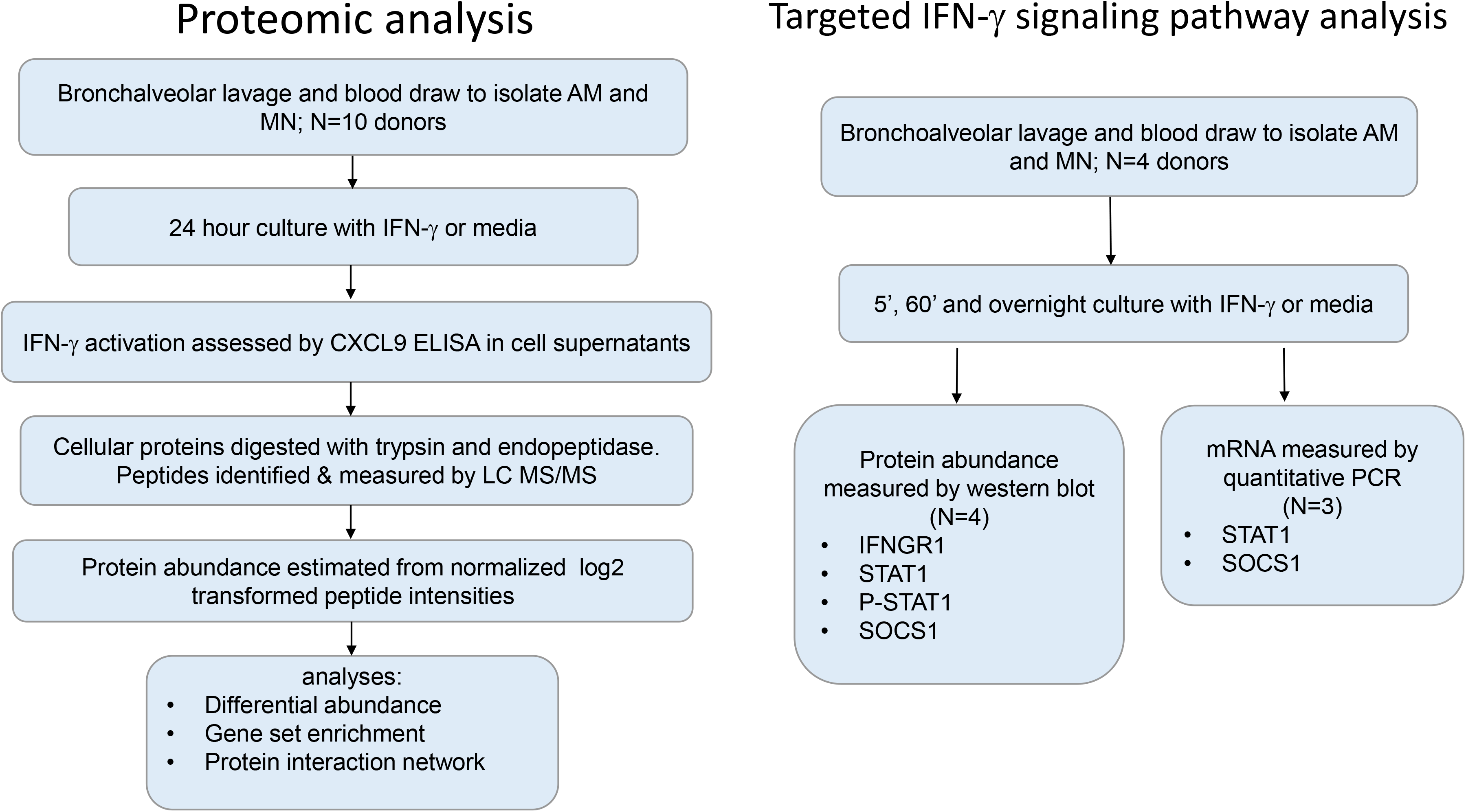
Workflow for exploratory proteomic analysis of autologous AM and MN samples stimulated with media or IFN-γ and the independent validation of STAT1 signaling and SOCS1 protein and gene expression in AM and MN.

Protein abundance was quantified by western blot band intensity relative to GAPDH band intensity. The relative intensity was assumed to be normally distributed and varying subject baselines were accounted for using the paired t-test comparing unstimulated and stimulated samples. Transcript mRNA abundance was measured by qPCR was normalized to GAPDH levels using the −2ΔCt method, and a paired t-test was used to test for a significant IFN-γ effect. Analyses were done in R (Version 4.0.4, Vienna, Austria). Results were graphed in Graphpad Prism for Windows V9.1.0 (San Diego, CA)

## RESULTS

### *Assessment of MIG (CXCL9) confirms* IFN-γ *-induced activation of both AM and MN*

For experiments evaluating the impact of IFN-γ on the AM and MN proteome, we utilized an IFN-γ concentration (1 U/ml;0.5 ng/ml) based on levels measured in the supernatants of similar cultures of BAL cells from individuals with latent tuberculosis infection (LTBI) following overnight incubation with PPD. IFN-γ-induced activation of AM and MN from healthy Mtb-naïve subjects was confirmed by measuring IFN-γ -dependent chemokine MIG (CXCL9) by ELISA in 24-hour culture supernatants. Significant increases in MIG production by both AM and MN for the 10 paired subject samples, used for proteomic studies, were measured in response to IFN-γ (Figure 2a; p=0.002) indicating that the AM cells were responsive to physiologic doses of IFN-γ.

**Figure 2:**
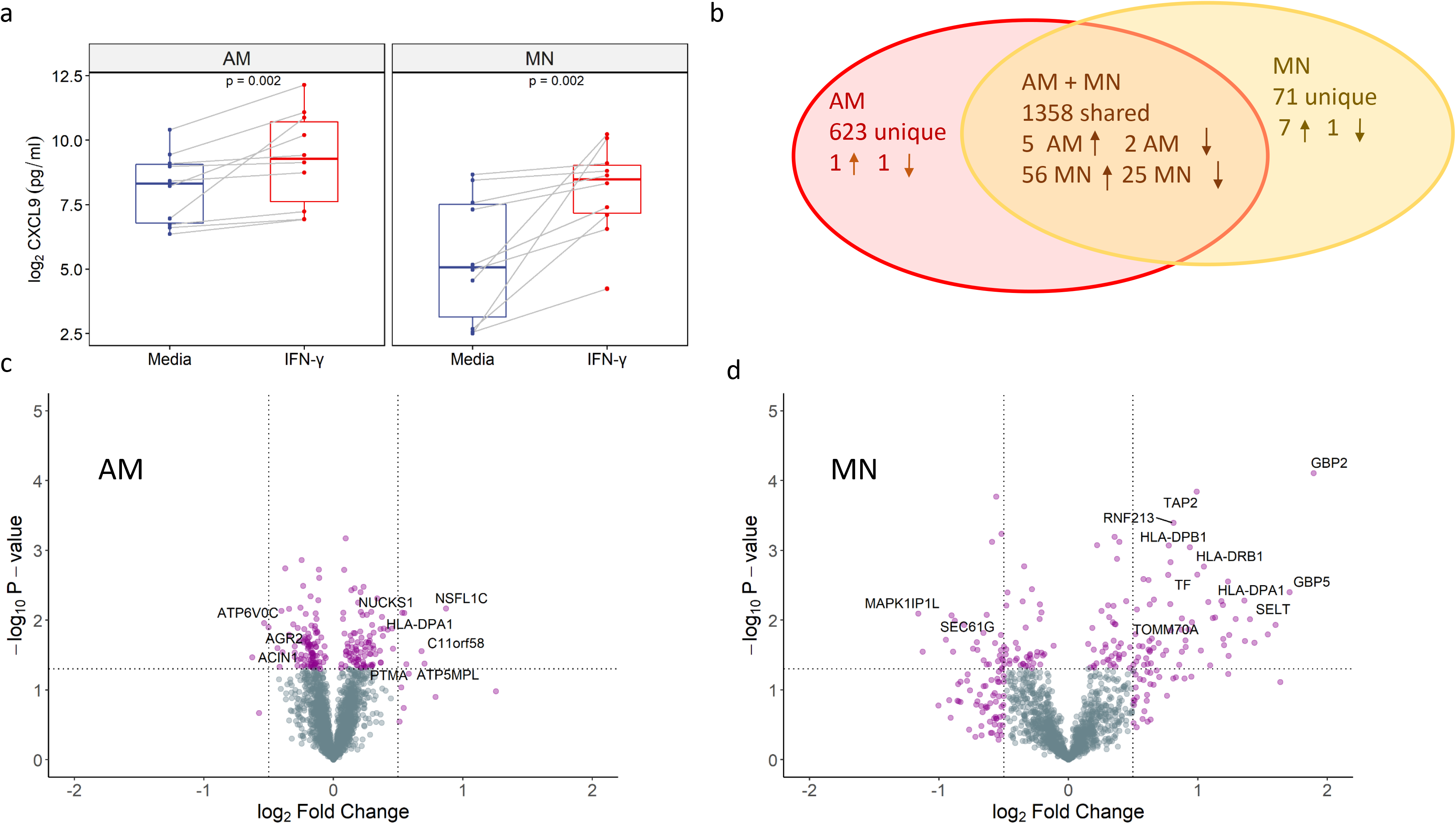
IFN-γ stimulation of AM and MN cells for 24 hours results in a larger number and percentage of differentially abundant proteins in MN relative to AM. a) CXCL9 (MIG) chemokine levels were measured in AM and MN culture supernatants by ELISA with and without 24 hr IFN-γ stimulation. Lines are shown connecting the paired results for each donor. b) Venn diagram with the number of proteins discovered within and across both cell types and the number of proteins up- and down-regulated based on *p*-value < 0.05 and absolute value fold change (FC) > 0.5. c) MN and d) AM volcano plots showing the significance (−log_10_ *p*-value) vs the log_2_ fold change in protein abundance. Data reflect the results from paired MN and AM from 10 healthy adult volunteers.

### *The AM proteomic response to IFN-*γ *is markedly less robust than that of MN*

The 10 matched donor AM and MN samples with and without IFN-γ stimulation were measured by LC MS/MS. Runs for all AM samples were aligned and aggregated such that there were no missing values, yielding 16,346 peptides mapped to 2,538 unique proteins under both stimulated and unstimulated conditions. Similarly, for the MN samples, 7,199 peptides were identified and mapped to 1,553 proteins. The median number of peptides discovered per protein in the combined AM and MN datasets was 6 (min=1, max=184). After filtering out proteins identified by only in a single peptide across both AM and MN datasets, there were 1981 AM and 1429 MN proteins. Figure 2b shows the number of proteins found within and across AM and MN including the number of significantly differentially abundant proteins.

Testing each protein for a significant IFN-γ treatment effect yielded 89 MN proteins and 9 AM proteins using a *p*-value < 0.05 and absolute fold change (|FC|) > 0.5 as significance criteria. Supplemental dataset File1 provides a list of all filtered proteins with estimated treatment effects and *p*-values. Of the 9 differentially abundant AM proteins, 6 increased in and 3 decreased in abundance, in contrast, in MN 63 proteins increased and 26 decreased in response to IFN-γ. Volcano plots for AM and MN proteins are shown in figures 2c and 2d. The 6 significantly increased proteins in AM were ATP5MPL, C11orf58, HLA-DPA1, NSFl1C, NUCKS1 and PTMA. The 3 significantly downregulated proteins were ACIN1, AGR2, and ATP6VOC. For MN the top proteins with increased abundance included GBP2, TAP2, HLA-DPB1, HLA-DPA1, HLA-DRB1 and STAT1. The most downregulated MN proteins included MAPK1IP1L, SEC61G and CALM5.

To identify functional relationships between proteins regulated by IFN-γ in each cell type, we tested for significant enrichment of MsigDB gene sets among all proteins ranked by the IFN-γ t-statistic. Figure 3a shows a comparison of gene sets enriched among AM and MN ranked proteins. The top enrichment in MN contained upregulated proteins involved in IFN-γ and cytokine responses whereas in AM these same gene sets were upregulated to a lesser extent and did not reach statistical significance for enrichment. The only gene sets found to be significantly enriched among AM proteins were downregulated and related to lipid modification and catabolic processes.

**Figure 3:**
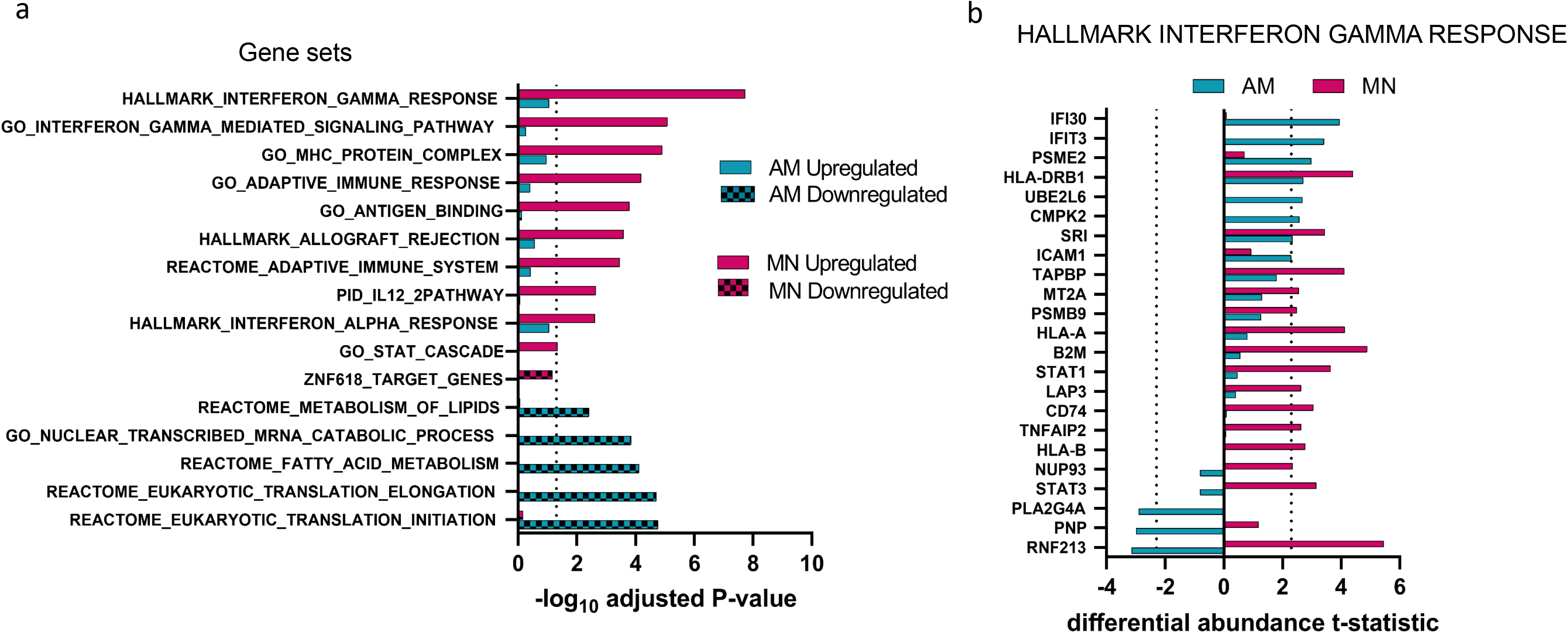
IFN-γ response signatures were significantly enriched with upregulated MN proteins in contrast to AM proteins which showed no significant upregulated enrichment. a) AM and MN proteins were ranked based on the IFN-γ differential abundance effect size and analyzed for signature enrichment. Dotted line at 1.3 delineates enrichment scores with adjusted *p*-value < 0.05. b) AM and MN proteins within the most significantly enriched gene set, *Hallmark Interferon Gamma Response*, illustrate that more proteins in this set are differentially regulated in MN (14) relative to AM (6 upregulated, 3 downregulated). The x-axis shows the differential abundance t-statistic for each protein with dotted lines at significance level = 0.05

To determine if proteins in canonical IFN-γ pathways were absent from AM cells or simply had lower abundance changes relative to MN, we focused on the proteins within the *Hallmark IFN-γ Response* gene set which is a curated collection of genes involved in IFN-γ regulated pathways and processes. As shown in figure 3b, many of the proteins that were upregulated in MN were found in AM but had smaller IFN-γ responses. In fact, several proteins in the canonical IFN-γ response were downregulated in AM. In summary, we expected IFN-γ to significantly alter the abundance of proteins involved in the classical IFN-γ regulated immune response but found that at a proteomic level, AM cells were minimally responsive. In contrast, MN were much more responsive to low levels of IFN-γ stimulation.

### Differences in the IFN-γ-induced proteomic responses of AM vs MN are not explained by alterations of IFN-γ signaling pathways

The diminished response in AM to IFN-γ relative to MN suggested the possibility that the proximal IFN-γ signaling pathway was inhibited or less activated in AM. We measured IFNGR1, P-STAT1, STAT1 and SOCS1 by western blot to look for differences in AM and MN protein levels over time. IFNGR1 levels were similar in AM and MN before and after IFN-γ stimulation (figure 4a). Figure 4b shows that STAT1 levels were constitutively higher in AM relative to MN and IFN-γ stimulation did not affect levels of STAT1 in AM. In contrast, STAT1 protein was significantly increased at the overnight time point in MN (p=0.02). In figure 4c, P-STAT1 was significantly induced by IFN-γ treatment at all time points in AM and was only increased in MN by the overnight time point (p=0.02). Levels of the SOCS1 repressor protein were low in both cell types and there were no constitutive differences or significant changes with IFN-γ stimulation (figure 4d).

**Figure 4:**
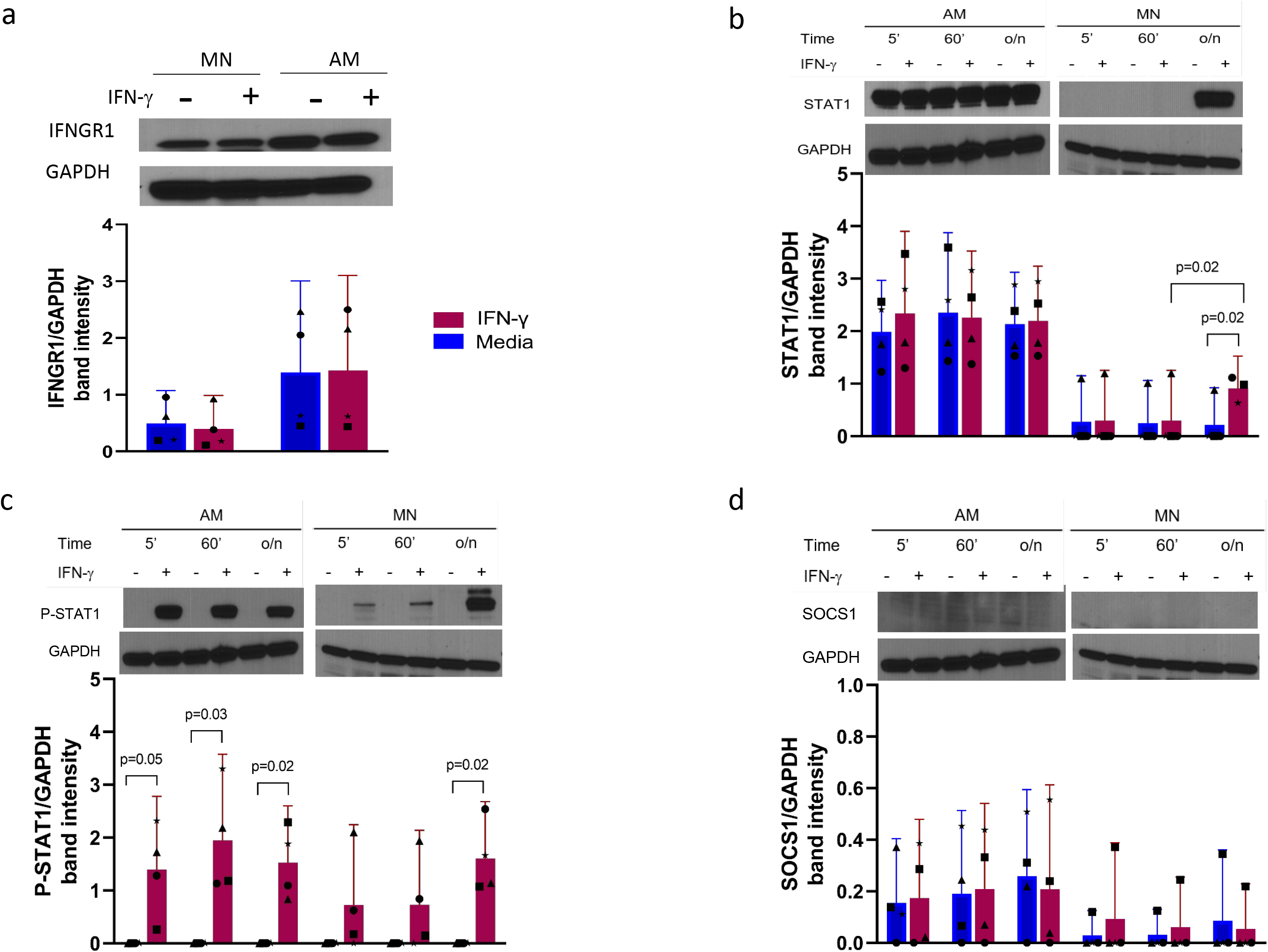
Differences in the proteomic outcomes of IFN-γ stimulation of AM vs MN are not associated with major functional proteins of the IFN-γ proximal signaling pathway. a) IFNGR1 protein in MN and AM with and without overnight (o/n) IFN-γ stimulation shows robust receptor levels that were not associated with treatment. b) STAT1/GAPDH relative protein abundance over time in AM and MN in the presence and absence of IFN-γ show abundant constitutive STAT1 levels in AM and up-regulation of STAT1 in MN following overnight treatment. c) P-STAT1 protein abundance in AM and MN show phosphorylation of STAT1 in response to IFN-γ and upregulation of in MN after overnight treatment. d) SOCS1 protein abundance time course in the presence and absence of IFN-γ showed low levels that were not associated with treatment. Band intensities were normalized to GAPDH levels (N=4). Representative western blots are shown above the plots. Bar plots are mean +/− 95% CI. Symbol shapes (●■▲*) represent a single donor.

We also measured STAT1 and SOCS1 transcript levels over time, with and without IFN-γ stimulation. Figure 5a shows that STAT1 mRNA from matched MN and AM from 3 donors at baseline and following 5 min, 1 hour and overnight incubation with IFN-γ resulted in no significant change in total STAT1 transcription in either cell type. SOCS1 mRNA levels are shown in figure 5b. Baseline levels of were low to undetectable in both AM and MN, and again there were no significant changes with IFN-γ stimulation in either cell type. The hypo-responsiveness in AM relative to MN is not due to a decrease in proximal signal transduction via JAK-STAT1 as assessed via both STAT1 transcription and STAT1 phosphorylation; nor is it due to an upregulation of the SOCS1 cytokine suppressor protein or SOCS1 transcript.

**Figure 5:**
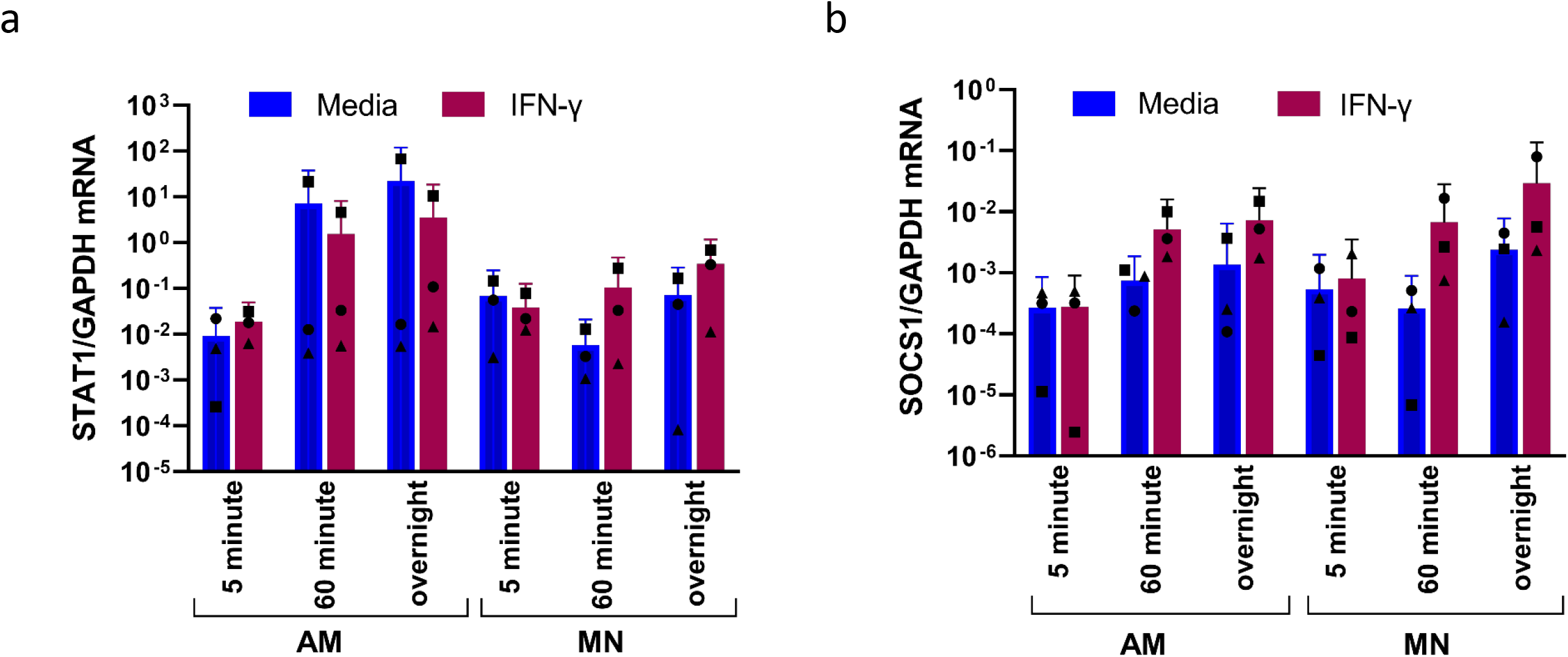
STAT1 and SOCS1 mRNA levels are not significantly affected by IFN-γ stimulation in AM and MN. a) STAT1 and b) SOCS1 transcript levels measured by quantitative PCR show no IFN-γ response differences after 5 minutes, 1 hour or overnight stimulation. Transcript levels were normalized to GAPDH (N=3). Bar plots are mean +/− 95% CI. Symbol shapes (●■▲) represent individual donors.

## DISCUSSION

We modeled the Th1 adaptive immune response in the lung at a proteomic level using ex-vivo stimulation of AM with physiologic doses of IFN-γ (0.5 ng/ml, or 1.0 unit/ml) and compared these responses to those of autologous blood MN. Although this dose was substantially less than used in many studies of IFN-γ dependent macrophage activation, secretion of CXCL9 chemokine measured by ELISA was similar in the two cell types indicating that both cell types responded to this treatment. Total proteomic abundance changes were measured using unlabeled mass spectrometry to provide a comprehensive view of the protein interactions and pathways in an overnight IFN-γ activation snapshot. At the proteomic level, a similar number of proteins were detected in both cell types. IFN-γ treatment had a significant effect on 9/1981 proteins in AM and 89/1429 in MN with larger abundance changes occurring in MN. Gene set enrichment analysis showed that proteins involved in the classic IFN-γ immune response were significantly enriched among the upregulated MN proteins showed little enrichment in AM. This effect may be due to an active down-regulated response in AM cells or the absence of the ability to mount a sustained inflammatory response.

Several gene sets showed significant enrichment of downregulated proteins in AM and no enrichment in MN. Downregulation of AM proteins in gene sets involved in lipid and fatty acid metabolism suggest a switch to glycolysis from oxidative phosphorylation to supply cellular energy, however this switch is generally associated with an increased inflammatory/M1 phenotype in macrophages that have been stimulated by LPS [10,11]. It is not clear whether a decrease in proteins related to the oxidative phosphorylation pathway is associated with a global reduction in the IFN-γ stimulated response and this an area of future investigation.

We found a relative resistance to Th1 cytokine stimulation in human AM despite increased CXCL9 secretion in both cell types. Since AM secrete CXCL9 in response to IFN-γ stimulation and CXCL9 is known to be expressed through JAK-STAT signal transduction [12], we focused on proteins involved in STAT1 signaling to test for differential function in AM and MN using an independent set of donors.

Binding of IFN-γ to the receptor starts a cascade that results in phosphorylated STAT1 translocation to the nucleus and transcription of interferon stimulated genes (ISGs) including the STAT1 gene itself. We found strong induction of P-STAT1 protein in AM where STAT1 protein levels were constitutively high at the 5-minute, 1 hour and overnight time points, but STAT1 levels did not change in response to IFN-γ treatment (Figures 4b and 4c). In contrast, STAT1 and P-STAT1 proteins were significantly upregulated in MN in response to IFN-γ at the overnight time point, suggesting that transcriptional activation and translation of STAT1 protein did occur in MN. Measurement of mRNA STAT1 levels in AM and MN did not show increased transcription in response to treatment (Figure 5a). We expected to see an increase in STAT1 transcript by the 60-minute time point and although a significant increase was not seen, figure 5a shows that all 3 donor samples had an increase in MN STAT1 mRNA in the treated condition. These results suggest that proximal signaling via the JAK-STAT pathway is intact in both AM and MN, but that subsequent transcription and translation was only upregulated in MN.

Next, we hypothesized that SOCS1 protein, which inhibits IFN-γ signaling by blocking JAK phosphorylation, might be constitutively high or induced by IFN-γ in AM cells. Comparison of mRNA and SOCS1 protein levels in treated and untreated conditions showed levels that were very low to undetectable. SOCS1 transcript and protein were not induced by IFN-γ in either AM or MN. Based on these results it is unlikely that SOCS1 is acting as a repressor of AM cell activation. In addition, the robust phosphorylation of STAT1 in AM does not support a role for SOCS1 inhibition.

Significant upregulation of a large number of proteins in AM cells was not necessarily expected given other studies of lung AM showing a tendency towards homeostasis following antigen stimulation. Previous studies have shown a suppressed response in murine AM where downregulation of bacterial clearance occurs with IFN-γ exposure [13,14]. Less is known about the human AM response to cytokine stimulation although several studies have shown evidence that AM macrophages are poor activators of T-lymphocytes relative to autologous MN [15] possibly due to lower induction of surface co-stimulatory molecules [16]. Lung macrophages in healthy individuals do not appear to be polarized towards either an inflammatory or anti-inflammatory phenotype. Recent studies indicate that fetal-derived AM of mesothelial origin populate the alveolar space during steady-state conditions, and while they have the capacity for self-renewal, they are augmented by circulating MN in response to lung injury and are replaced by MN during healthy aging [17,18]. Therefore, the refractory phenotype in AM in response to cytokine could be due to the embryonic cell origin as distinct from bone marrow derived MN.

Our interest in IFN-γ-responsiveness of MN and AM developed from our BAL cell findings in individuals with LTBI, for whom IFN-γ-induced responses provide the dominant component of an Mtb-induced local recall gene expression signature [19]. Further studies have indicated that both peripheral blood and BAL cells in LTBI uniquely display baseline expression of an immune response gene signature that is not observed in AM of Mtb-BCG-naïve controls or of recipients of intradermal or oral BCG vaccine (Silver, submitted). These findings could indicate epigenetic AM conditioning in LTBI that could manifest as increased responsiveness of AM in LTBI to IFN-γ stimulation. Given that studies in LTBI subjects were the basis for our choice of IFN-γ dosing in the current study, it is possible that higher concentrations of IFN-γ may be required to induce more robust responses by AM from the Mtb-naïve subjects as studied here. On the other hand, the current use of low-dose IFN-γ in our studies allowed for the demonstration of substantial differences in responses of MN and AM from Mtb-naïve human subjects to physiologic concentrations of IFN-γ.

This study had several limitations. LC MS/MS can measure thousands of peptides, however, there is a limitation on measurement of low abundance and secreted proteins and therefore chemokines such as CXCL9, expected to be regulated by IFN-γ, could not be detected in the proteomics assay. In addition, the proteomics snapshot at 24 hours may have missed some of the protein signaling occurring earlier or later in AM cells.

Previous studies have shown a reduced innate inflammatory response of human AM to infection or antigen stimulation. In this study we modeled T-cell derived IFN-γ stimulation of AM and demonstrated a proteomic hypo-responsiveness relative to circulating MN that is not due to a reduction in the proximal signaling pathway. These findings may serve as a basis for investigating whether AM in LTBI and other conditions of prolonged antigenic stimulation may result in epigenetic conditioning resulting in increased AM responsiveness to IFN-γ stimulation.

## ACKNOWLEGEMENTS

We thank the healthy volunteers who underwent research BAL and blood draws for this study, as well as the nurses and staff of the Dahms Clinical Research Unit who supported this work.

## REFERENCES

[1] B. Allard, A. Panariti, J.G. Martin, Alveolar Macrophages in the Resolution of Inflammation, Tissue Repair, and Tolerance to Infection, Front Immunol. 9 (2018) 1–7. https://doi.org/10.3389/fimmu.2018.01777.

[2] B.M.N. Kagina, B. Abel, T.J. Scriba, E.J. Hughes, A. Keyser, A. Soares, H. Gamieldien, M. Sidibana, M. Hatherill, S. Gelderbloem, H. Mahomed, A. Hawkridge, G. Hussey, G. Kaplan, W.A. Hanekom, Specific T cell frequency and cytokine expression profile do not correlate with protection against tuberculosis after bacillus Calmette-Guérin vaccination of newborns, Am J Respir Crit Care Med. 182 (2010) 1073–1079. https://doi.org/10.1164/rccm.201003-0334OC.

[3] M.D. Tameris, M. Hatherill, B.S. Landry, T.J. Scriba, M.A. Snowden, S. Lockhart, J.E. Shea, J.B. Mcclain, G.D. Hussey, W.A. Hanekom, H. Mahomed, H. Mcshane, Safety and efficacy of MVA85A, a new tuberculosis vaccine, in infants previously vaccinated with BCG: a randomised, placebo-controlled phase 2b trial, The Lancet. 381 (2013) 1021–1028. https://doi.org/10.1016/S0140-6736(13)60177-4.

[4] S.E. Tomechko, K.C. Lundberg, J. Jarvela, G. Bebek, N.G. Chesnokov, D. Schlatzer, R.M. Ewing, W.H. Boom, M.R. Chance, R.F. Silver, Proteomic and bioinformatics profile of paired human alveolar macrophages and peripheral blood monocytes, Proteomics. 15 (2015) 3797–3805. https://doi.org/10.1002/pmic.201400496.

[5] J. Walrath, L. Zukowski, A. Krywiak, R.F. Silver, Resident Th1-like effector memory cells in pulmonary recall responses to Mycobacterium tuberculosis, Am J Respir Cell Mol Biol. 33 (2005) 48–55. https://doi.org/10.1165/rcmb.2005-0060OC.

[6] M. Choi, C.-Y. Chang, T. Clough, D. Broudy, T. Killeen, B. MacLean, O. Vitek, MSstats: an R package for statistical analysis of quantitative mass spectrometry-based proteomic experiments, Bioinformatics. 30 (2014) 2524–2526. https://doi.org/10.1093/bioinformatics/btu305.

[7] T. Clough, S. Thaminy, S. Ragg, R. Aebersold, O. Vitek, Statistical protein quantification and significance analysis in label-free LC-MS experiments with complex designs., BMC Bioinformatics. 13 Suppl 1 (2012) S6. https://doi.org/10.1186/1471-2105-13-S16-S6.

[8] A. Liberzon, A. Subramanian, R. Pinchback, H. Thorvaldsdóttir, P. Tamayo, J.P. Mesirov, Molecular signatures database (MSigDB) 3.0, Bioinformatics. 27 (2011) 1739–1740. https://doi.org/10.1093/bioinformatics/btr260.

[9] A. Liberzon, C. Birger, H. Thorvaldsdóttir, M. Ghandi, J.P. Mesirov, P. Tamayo, The Molecular Signatures Database Hallmark Gene Set Collection, Cell Syst. 1 (2015) 417–425. https://doi.org/10.1016/j.cels.2015.12.004.

[10] P.S. Woods, L.M. Kimmig, A.Y. Meliton, K.A. Sun, Y. Tian, E.M. O’Leary, G.A. Gökalp, R.B. Hamanaka, G.M. Mutlu, Tissue-resident alveolar macrophages do not rely on glycolysis for LPS-induced inflammation, Am J Respir Cell Mol Biol. 62 (2020) 243–255. https://doi.org/10.1165/rcmb.2019-0244OC.

[11] J. van den Bossche, L.A. O’Neill, D. Menon, Macrophage Immunometabolism: Where Are We (Going)?, Trends Immunol. 38 (2017) 395–406. https://doi.org/10.1016/j.it.2017.03.001.

[12] K. Schroder, P.J. Hertzog, T. Ravasi, D.A. Hume, Interferon-y◻: an overview of signals, mechanisms and functions, J Leukoc Biol. 75 (2004) 163–189. https://doi.org/10.1189/jlb.0603252.Journal.

[13] K. Sun, D.W. Metzger, Inhibition of pulmonary antibacterial defense by interferon-γ during recovery from influenza infection, Nat Med. 14 (2008) 558–564. https://doi.org/10.1038/nm1765.

[14] M.J. Mina, L.A.S. Brown, K.P. Klugman, Dynamics of Increasing IFN-γ Exposure on Murine MH-S Cell-Line Alveolar Macrophage Phagocytosis of Streptococcus pneumoniae, Journal of Interferon and Cytokine Research. 35 (2015) 474–479. https://doi.org/10.1089/jir.2014.0087.

[15] P.G. Holt, Down-regulation of immune responses in the lower respiratory tract: The role of alveolar macrophages, Clin Exp Immunol. 63 (1986) 261–270.

[16] C.J. Chelen, Y. Fang, G.J. Freeman, H. Secrist, J.D. Marshall, P.T. Hwang, L.R. Frankel, R.H. Dekruyff, D.T. Umetsu, Human Alveolar Macrophages Present Antigen Ineffectively due to Defective, Journal of Clinical Investigation. 95 (1995) 1415–1421.

[17] A.J. Byrne, J.E. Powell, B.J.O. Sullivan, P.P. Ogger, A. Hoffland, J. Cook, K.L. Bonner, R.J. Hewitt, S. Wolf, P. Ghai, S.A. Walker, S.W. Lukowski, P.L. Molyneaux, S. Saglani, D.C. Chambers, T.M. Maher, C.M. Lloyd, Dynamics of human monocytes and airway macrophages during healthy aging and after transplant, Journal of Experimental Medicine. 217 (2020) 1–11.

[18] E. Evren, E. Ringqvist, K.P. Tripathi, N. Sleiers, I. Co Rives, A. Alisjahabana, Y. Gao, D. Sarhan, T. Halle, C. Sorini, R. Lepzien, N. Marquardt, J. Michaelsson, A. Smed-Sorenson, J. Botling, M.C.I. Karlsson, E.J. Villablanca, T. Willinger, J. Botling, M.C.I. Karlsson, E.J. Villablanca, T. Willinger, Distinct developmental pathways from blood monocytes generate human lung macrophage diversity ll ll Distinct developmental pathways from blood monocytes generate human lung macrophage diversity, Immunity. 54 (2021) 259–275. https://doi.org/10.1016/j.immuni.2020.12.003.

[19] J. Jarvela, M. Moyer, P. Leahy, T. Bonfield, D. Fletcher, W.N. Mkono, H. Aung, D.H. Canaday, J.-E. Dazard, R.F. Silver, Mycobacterium tuberculosis –Induced Bronchoalveolar Lavage Gene Expression Signature in Latent Tuberculosis Infection Is Dominated by Pleiotropic Effects of CD4 + T Cell–Dependent IFN-γ Production despite the Presence of Polyfunctional T Cells within the Airways, The Journal of Immunology. 203 (2019) 2194–2209. https://doi.org/10.4049/jimmunol.1900230.

